# Reduction of Deformed Wing Virus-B levels in Colonies of the Honey bee *Apis mellifera* after Queen Vaccination with inactivated *Paenibacillus larvae*

**DOI:** 10.1101/2024.07.01.601551

**Authors:** Isaac Weinberg, Amy Floyd, Nathan Reid, Nigel Swift

## Abstract

Deformed wing virus (DWV) is a virulent and ubiquitous disease that affects honey bee colonies. DWV infects bees at all life stages, but is most noticeable in adult bees, where clinical symptoms include shriveled, non-functional wings, and a drastically shortened lifespan. DWV is recorded in upwards of 90% of honey bee colonies worldwide, and has been linked to colony loss in symptomatic hives. DWV is primarily spread by *Varroa destructor* mites, who feed on the fat bodies of honey bee adults and pupae. There is currently no direct treatment or preventative for DWV, with the primary method of reduction being vector control using acaracides. In this study, we tested the effect that vaccinating honey bee queens using killed *Paenibacillus larvae* bacterin, the causative agent of the honey bee disease American Foulbrood, had on deformed wing virus load in honey bee colonies. We placed vaccinated queens in 200 honey bee colonies, and unvaccinated queens in 200 colonies, and measured quantities of DWV-B in both groups immediately before, and 4 months after vaccination. We found that levels of DWV-B were identical before vaccination, but were significantly reduced in colonies 4 months post vaccination. This change was found despite no difference in mite quantities between groups. Overall, these data provide evidence that vaccination of queens with *P. larvae* bacterin is an effective method for reduction of DWV-B quantities in honey bee colonies in a commercially relevant field setting.

## Introduction

Deformed Wing Virus (DWV), an RNA virus in the family Iflaviridae, is one of the most widespread and destructive diseases affecting honey bees (Chen & Siede, 2007; de Miranda & Genersch, 2010), with studies commonly finding DWV in over 90% of tested colonies (Natsopoulou et al., 2017; Kevill et al., 2019; Paxton et al., 2022). Pupae infected with DWV develop into abnormal adult bees with undeveloped non-functional wings, bloated abdomens, decreased adult size, and severely reduced lifespan (de Miranda & Genersch, 2010). There is evidence that even subclinical levels of DWV in bees can impair cognitive function, reduce foraging efficacy, and reduce lifespan (Benaets et al., 2017; Chen et al., 2021).

DWV is split into 3 main strains: the common DWV-A (Lanzi et al., 2006) and DWV-B (Ongus et al., 2004), and the less common DWV-C (Mordecai et al., 2016). While DWV-A has historically been most prevalent, DWV-B is rapidly displacing it worldwide in association with the global spread of *Varroa destructor* mites. (Paxton et al., 2022). DWV-B accumulates at higher levels in bees and brood (Norton et al., 2020) potentially because it is more easily transmitted by *Varroa destructor* than DWV-A (Ryabov et al., 2014, 2019). As a result of their extensive pathology, both DWV-A and DWV-B have been directly linked to overwintering failure (Kevill et al., 2019; Natsopoulou et al., 2017), and are of tremendous concern to beekeepers.

DWV is highly associated with the obligate honey bee ectoparasite mite, *Varroa destructor* (Wilfert et al., 2016; Barroso-Arévalo et al., 2019; Posada-Florez et al., 2019) the factor beekeepers most strongly associate with colony loss (Engebretson et al., 2022). *Varroa* mites parasitically feed on the fat body tissue of both developing pupae and adult bees (Ramsey et al., 2019; Warner et al., 2024) making them an effective vector for DWV transmission (Natsopoulou et al., 2017; Posada-Florez et al., 2019). *Varroa* mites also increase the severity of DWV infection by weakening their host’s immune system through down regulation of host immune gene expression (Nazzi et al., 2012). In the absences of *Varroa*, DWV persists in colonies through vertical transmission from infected queens to their offspring, or by direct horizontal transmission via trophallaxis or larval feeding (Yue & Genersch, 2005; Yue et al., 2007). Typically, miteless transmission produces low viral loads, and results in subclinical DWV (Locke et al., 2017).

Studies have shown that activating the RNAi-system by feeding virus-specific dsRNA to larvae or adult bees before infection with DWV or Israeli Acute Paralysis Virus (IAPV) reduces the viral load, mortality, and symptoms resulting from the specific viral infection (Hunter et al., 2010; Desai et al., 2012). Potassium ion channel manipulation has also been proven effective treatment for IAPV (Fellows et al., 2023). Neither of these methods, however, are currently approved for use in the field, and there is no licensed specific treatment or prophylaxis for DWV (Smeele et al., 2023). Currently, the primary method of reducing DWV levels in a colony is vector control through the reduction of *Varroa* levels (Woodford et al., 2022). This is generally done using insecticides and acaricides like amitraz (Warner et al., 2024) which are widely used in agriculture for pest control. However, when used in honey bees, these chemicals can reduce viral immunity (O’Neal et al., 2017) and contaminate honey (Pohorecka et al., 2018).

An alternate mechanism for direct disease management in honey bee colonies is protection through vaccination. Currently, vaccination using the Paenabicillus larvae bacterin is commercially approved for protection of honey bee larvae against American Foulbrood, through Trans Generational Immune Priming (TGIP), the parent-to-offspring transfer of immune experience (Hernández López et al., 2014; Dickel et al., 2022). Vaccinating parent insects with killed pathogens has been shown to provide homologous protection through TGIP across a wide range of invertebrate lineages (Tetreau et al., 2019) including honey bees (Freitak et al., 2014; Hernández López et al., 2014). It has been hypothesized that vaccination using inactivated DWV-A could provide trans-generational protection, though results to date are mixed, with context-dependent TGIP benefits recorded after venereal transmission of DWV-A to queen bees (Lang et al., 2022), but a lack of benefits after oral administration (Leponiemi et al., 2021).

Here, we present the first evidence that oral vaccination of a queen bee with an inactivated gram-positive bacteria, *Paenibacillus larvae* can directly reduce DWV-B levels in honey bee colonies in the field, independent of *Varroa destructor* burden. These data are to our knowledge the first use of TGIP with a killed bacteria to provide heterologous protection against a virus, and the first example of control of DWV in the field independent of mite control.

## Methods

### Vaccination of Queens

On May 31, 2023, 400 Italian honey bee queens, *Apis mellifera ligustica*, were sourced from Vidalia Apicultural Services in Toombs County GA. 200 queens were vaccinated with fully inactivated *P. larvae* bacterin (PCN 2915.00, produced according to the proprietary outline of production of Diamond Animal Health) and 200 remained unvaccinated. For every 50 bees to be vaccinated, 3mL of *P. larvae* bacterin solution was added to 300g of queen candy (approx. 1:7.5 w/w corn syrup to powdered sugar), resulting in 1.5*10^8^ dead *P. larvae* cells per queen. Queens were isolated in queen cages and placed battery boxes in groups of 50. 300g of candy containing the bacterin was laid out in two strips between the queen cages in each battery box. Approximately 2100 nurse bees were then placed in the battery box to attend to the queens. Queens and nurses were left in the battery boxes for 8 days to allow for total consumption of the candy, before being placed into hives. Due to difficulties in procuring 400 queens on the same day, the 200 unvaccinated queens did not go through the vaccination protocol, and are therefore true controls and not placebo controls. Requeening of all hives occurred on June 8, 2023.

### Subjects

In May 2023, 400 established honey bee colonies across 8 yards in Toombs County Georgia were entered into the study. Yards contained between 39 and 78 colonies, which were divided roughly evenly into vaccinated and unvaccinated. Prior to study onset, colonies were inspected to verify they were robust and free of pests and disease before entry into the trial. At the start of the study, each colony consisted of one deep and one medium hive body. Most colonies maintained this size through the study, however some expansions and contractions were performed by yard crews in line with their standard hive management practices. Colonies were treated for *Varroa* mites monthly using amitraz, and were fed sugar water during periods of low nectar flow to supplement their nectar intake.

### Sample Collection

10 nurse bees were collected from 100 vaccinated and 100 unvaccinated colonies immediately before experimental queens were placed into colonies in May 2023, and then again 4 months after queens were placed in colonies in September 2023. Nurse bees were collected from above the brood comb, and were visually identified by their coloration and intact dorsal hairs. Nurse bees were placed into 15ml falcon tubes and flash frozen on dry ice in the field. Bees were then stored in a -80C freezer in the lab until submission.

### Sample Analysis

In all 8 yards, a single pooled sample of nurse bees was sent for analysis from each experimental group. Each pooled sample contained 50 bees collected equally from 10 colonies (N=8 samples/ treatment/ timepoint).

Blinded samples were sent to and analyzed by the National Agricultural Genotyping Center (NAGC) in Fargo SD, an accredited testing facility used by beekeepers and scientists across North America. DWV-B levels were analyzed using qPCR. Samples were tested against a positive control, reagent control, and a no template control. PCR cycling conditions are proprietary to the NAGC and therefore cannot be reported here.

Samples were tested using similar methods for DWV-A, DWV-C, American Foulbrood, European Foulbrood, and Sacbrood Virus, but little to no instances of these disease were observed, and those results are therefore not reported.

### Mite Quantification

Mites were collected from colonies both in May and November 2023. Counts were taken using a Veto-pharma *Varroa* Easy Check kit. Approximately 300 bees were collected from above the brood comb in sampled hives and washed in 70% ethanol to separate the mites from the bees. The quantity of mites beneath the sieve were quantified, and are reported as the number of mites/100 bees. Washes were taken from colonies In May (N=35 unvaccinated, N=38 vaccinated) and in November (N=44 vaccinated and N=44 unvaccinated).

### Statistical Analysis

DWV-B levels were compared using a Wilcox rank sum test because they were exponentially distributed. Mite counts were compared using a welch two sample T-test. All statistical analyses were conducted in R version 4.3.0 (R Core Team, 2024).

## Results

### DWV-B levels

DWV-B levels were identical between groups one week prior to vaccination (Vaccinated Median (IQR): 1.4*10^9^ (5.2*10^7^-2*10^9^); Unvaccinated Median (IQR): 3.6*10^8^ (2.6*10^8^-1.5*10^9^); Wilcoxon rank sum test, W=31, P=0.96), and were significantly reduced in vaccinated hives compared to control hives 4 months post vaccination (Vaccinated Median (IQR): 7.8*10^7^ (1.0*10^7^-2.2*10^8^); Unvaccinated Median (IQR): 1.4*10^9^ (7.9*10^8^-2.5*10^9^); Wilcoxon rank sum test, W=54.5, P=0.021) (Figure 1). DWV-B quantities were reduced in vaccinated colonies compared to unvaccinated colonies in all 8 yards, with an average reduction of 83%.

**Figure 1:**
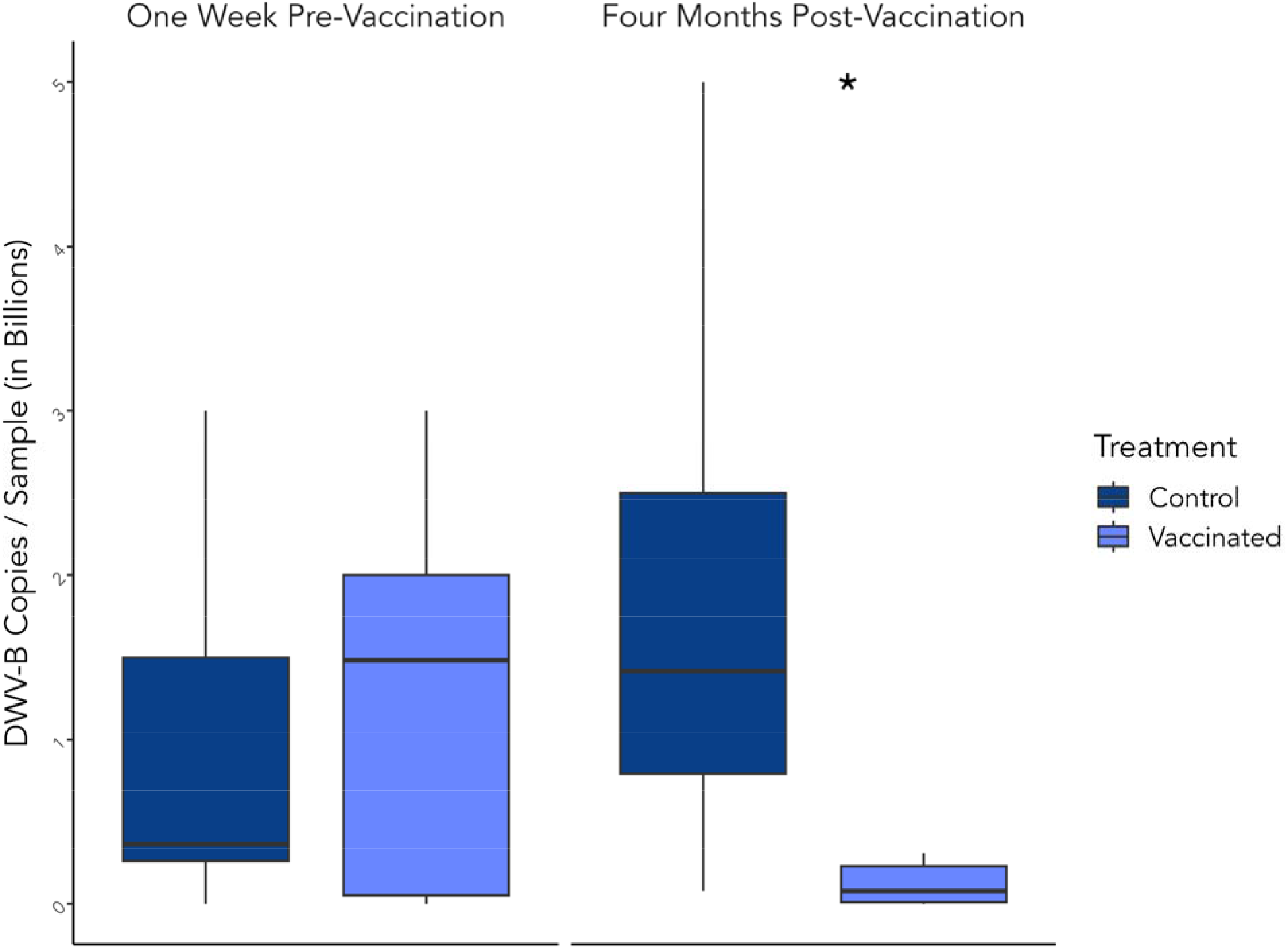
Box and whisker plot showing DWV-B copies per sample in vaccinated and unvaccinated honey bee colonies one week before and four months after vaccination. One sample from each experimental group was sent from each of the 8 yards at both timepoints (N=8/treatment/timepoint). Samples contained 50 bees pooled equally from 10 colonies in the same yard. Significance (P<0.05) on a Wilcox rank sum test is denoted by (*) between box and whisker plots.

### Mite Counts

There was no difference in mite counts between treatments one week prior to vaccination (Vaccinated Mean(SD): 1.9 (2.3) ; Unvaccinated Mean(SD): 2.1 (3.2); Welch two sample t-test, T=0.38, df=61 P= 0.71) or six months post vaccination (Vaccinated Mean(SD): 3.9 (10.4); Unvaccinated Mean(SD): 1.8 (2.4); Welch two sample t-test, T=-1.30, df=48 P= 0.20) (Figure 2). Following standard beekeeping practice, all colonies were regularly treated for mites using amitraz. These results indicate there was no difference in mite load between treatment groups.

**Figure 2:**
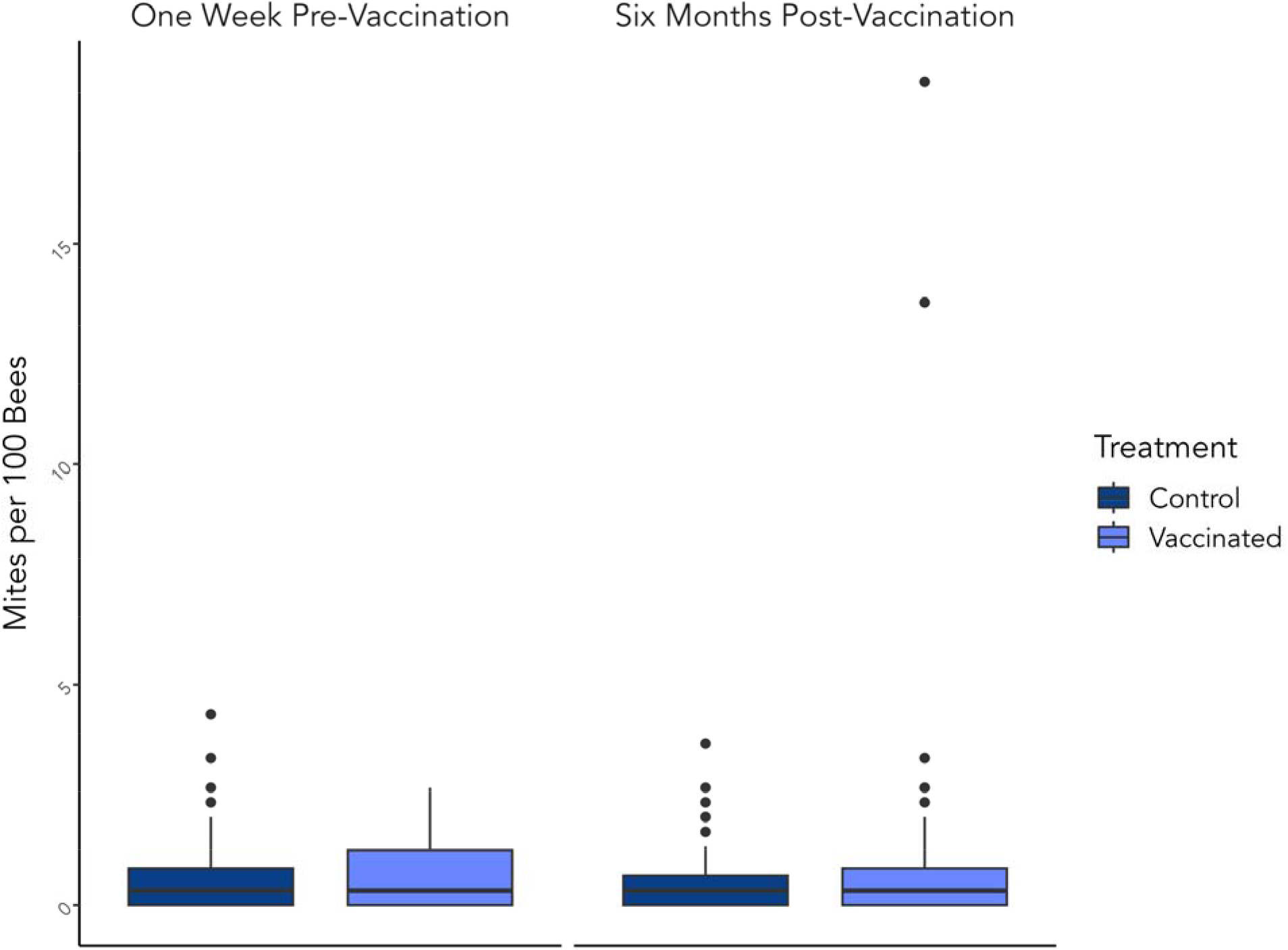
Box and whisker plot showing mite count in vaccinated and unvaccinated colonies one week before (N=35 unvaccinated, N=38 vaccinated) and six months after (N=44 vaccinated and N=44 unvaccinated) vaccination. Mite counts were taken by performing a standard alcohol mite wash.

## Discussion

This study provides the first evidence that vaccinating honey bee queens with Penibacillus larvae bacterin provides heterologous trans-generational protection against DWV-B in a field setting. These data are also to our knowledge the first evidence that vaccinating an organism with killed bacteria can provide protection against a viral disease in an organism possessing only an innate immune system. The reduction in viral load occurred despite there being no difference in *Varroa destructor* quantities between vaccinated and unvaccinated colonies, suggesting the effect is independent of mite load. This experiment was conducted on full size honey bee colonies at a commercial apiary, and involved no special colony management practices outside of providing colonies with vaccinated queens. These results indicate that vaccinating queens with the *P. larvae* bacterin may be an efficient method to directly control DWV-B in honey bee colonies in the field.

Most laboratory studies on DWV are undertaken in the absence of *Varroa destructor*, which are present in honey bee colonies worldwide, nearly impossible to eradicate, highly associated with increased DWV levels, and directly weaken and disrupt bees’ immune response (Annoscia et al., 2019; Kuster et al., 2014; Ramsey et al., 2019). Previous studies investigating homologous transgenerational protection against DWV in the lab have either failed to show protection (Leponiemi et al., 2021), or showed protection only under specific circumstances (Lang et al., 2022). That we observed heterologous protection against DWV-B after vaccinating queen bees with the *P. larvae* bacterin was therefore surprising. Arthropods, including honey bees, do not possess an adaptive immune system (Baxter et al., 2017) and rely solely on innate immunity for protection against pathogens (Morfin et al., 2021). It has been noted that honey bees express only 1/3 as many immune-regulating genes as solitary insects, and instead rely more on social immunity such as grooming, task separation and thermoregulation. This reduction in genes may increases “cross-talk” between immunologic pathways, and perhaps results in less specific immunologic responses which would increase the likelihood vaccination causes a generalized immune benefit.

While we have not yet determined the mechanism by which bacterial inoculation helps reduce viral infection, we speculate that the vaccine may be augmenting one or more of the common pathways of the innate bee immune system, thereby causing an increase in innate immune function of bee larvae (Nazzi & Le Conte, 2016; Annoscia et al., 2019). Innate immune responses often function by increasing expression of generalized defenses like AMPs and siRNAs (Kingsolver et al., 2013; Mondotte et al., 2020). It has been shown that bumble bee queens injected with heat-killed bacteria pass trans-generational immune benefits to their daughters, who upregulate genes associated with AMPs and toll signaling pathways, even in the absence of infection (Barribeau et al., 2016). Simply altering immune function, however, does not guarantee a decrease in disease. An upregulation of both RNAi genes and siRNA activity are observed in response to DWV infection, which does not inhibit DWV from accumulating to high levels (Norton et al., 2023). This may be in part due to an immunosuppression syndrome associated with DWV infection, which includes strong down-regulation of NF-kB, a transcription factor which helps protect against a wide range of environmental challenges (Nazzi et al., 2012). Future transcriptomic studies should look for modifications honey bee worker gene regulation after queens are inoculated with the *P. larvae* bacterin to shed light on how these complex interactions may be affecting bee immunity.

If the *P. larvae* bacterin does indeed improve general immune function, we would expect to see similar heterologous protection against a wide range of disease in the field. While we tested for protection against 5 other diseases, none were present at high enough levels in control or test groups to provide informative data. Future studies should continue to investigate cross-protective effects against other important honey bee disease like DWV-A, Chalkbrood, Israeli Acute Paralysis Virus, Sacbrood Virus, and Nosema. In conducting a field evaluation on the effects of vaccinating honey bee queens with *P. larvae* bacterin in a commercial apiary, we have demonstrated a significant reduction of naturally acquired DWV-B in honey bee colonies. DWV-B is one of the most detrimental diseases of honey bees, and this vaccine demonstrates potential to control DWV-B independent of mite treatment.

## Acknowledgements

We would like to thank Diamond Animal Health for manufacturing the *P. larvae* bacterin used in this study. We would also like to thank Vidalia Apicultural Services for providing the queens and colonies used in this study, and for performing colony maintenance. We would also like to thank Adam Costello, Jessica Lomasney, Kadijah Fennell, and Zoe York for help collecting data in the field. Finally, we would like to thank the National Agricultural Genotyping Center for conducing PCR analysis on our samples.

